# Variations and expression features of CYP2D6 contribute to schizophrenia risk

**DOI:** 10.1101/659102

**Authors:** Liang Ma, Sundari Chetty

## Abstract

**Objective:** Genome-wide association studies (GWAS) have successfully identified 145 loci implicated in schizophrenia (SCZ). However, the underlying mechanisms remain largely unknown.

**Methods:** Here, we analyze 1,479 RNA-seq data from 13 postmortem brain regions in combination with their genotype data and identify SNPs that are associated with expression throughout the genome by dissecting expression features to genes (eGene) and exon-exon junctions (eJunction). Then, we co-localize eGene and eJunction with SCZ GWAS using SMR and mapping. Multiple ChIP-seq data and DNA methylation data generated from brain were used for identifying the causative variants. Finally, we used a hypothesis-free (no SCZ risk loci considered) enrichment analysis to determine implicated pathways.

**Results:** We identified 171 genes and eight splicing junctions located within four genes (*SNX19, ARL6IP4, APOPT1* and *CYP2D6*) that potentially contribute to SCZ susceptibility. Among the genes, *CYP2D6* is significantly associated with SCZ SNPs in both eGene and eJunction across the 13 brain regions. In-depth examination of the *CYP2D6* region revealed that a non-synonymous single nucleotide variant (SNV) rs16947 is strongly associated with a higher abundance of *CYP2D6* exon 3 skipping junctions. While we found rs133377 and two other SNPs in high linkage disequilibrium (LD) with rs16947 (*r*^2^ = 0.9539), histone acetylation analysis showed they are located within active transcription start sites. Furthermore, our data-driven enrichment analysis showed CYP2D6 is significantly involved in drug metabolism of tamoxifen, codeine and citalopram.

**Conclusions:** Our study facilitates an understanding of the genetic architecture of SCZ and provides new drug targets.

## Introduction

Schizophrenia (SCZ) is a debilitating, highly heritable and polygenic psychiatric condition affecting roughly 1% of the population. Recent GWAS have successfully identified 145 risk loci by meta-analysis of samples through the UK CLOZUK and the Psychiatric Genomics Consortium (PGC2) and compared allele frequency differences between SCZ patients and normal controls (1). While these findings have identified regions in the genome harboring SCZ genes, almost all of the regions include multiple genes that are located within the same recombination hotspot intervals, making it challenging to identify causal genes.

One essential approach to determining the function of the identified SNPs is to combine GWASs with gene expression in post-mortem human brains. Recently, several studies have applied these strategies to the 108 loci identified by the PGC in 2014 (2). Using genotyped RNA sequencing (RNA-seq) data generated by the CommonMind Consortium from postmortem dorsolateral prefrontal cortex (DLPFC), splicing Quantitative Trait Loci (QTL) were found to be signifixscantly enriched in SCZ risk loci (3). The enrichment of SCZ loci was also observed in another independent study of postmortem brain DLPFC samples (4), suggesting that combining these analytical techniques provides important insights into the mechanisms underlying SCZ.

Studies integrating expression QTL (eQTL) and GWAS data have almost exclusively used quantified expression across multiple transcript features, including annotated genes as well as annotation-guided transcripts. Notably, junction calls from short RNA-seq reads are considerably more reliable than assembled transcripts (5). However, only a limited number of studies employed exon-exon splice junctions to identify splicing transcripts that contribute to the underlying phenotypes (4). Recently, skipping of exon 2 and exon 3 of AS3MT was identified to contribute to SCZ risk using large postmortem brain cohorts (6). Using the same specimens, we also successfully identified risk allele of SCZ SNPs are strongly associated with splicing junctions between exon 8 and exon 10 of *SNX19* (7).

Here, we leveraged genotype and brain expression data provided by the Genotype-Tissue Expression (GTEx) project to elucidate the functional properties and potential roles of eQTL and splicing expression features in the etiology of SCZ. Integrating the GTEx brain eQTLs with the most recent SCZ GWAS data (CLOZUK +PGC2) (1), we evaluated specific effects of the identified SCZ risk variants on gene expression features in combination with epigenetics data, and examined how genes of expression features affect genetic networks. Our co-localization analyses identify specific targets implicated in SCZ and highlight new biological insights for further investigation in future studies.

## Materials and methods

### RNA-seq of postmortem brain

A total of 1,497 human brain samples across 13 brain regions were used in this study (Table 1). All of the postmortem brain samples were collected by the GTEx consortium. The sample procurement has been described previously (8). Raw gene and exon-exon junction reads counts were retrieved from GTEx portal (https://gtexportal.org/home/datasets). Gene lengths were calculated using GENCODE v19 annotations (9). We converted gene counts to RPKM (Reads per Kilobase per Million mapped reads) values using the total number of aligned reads across the 22 autosomal chromosomes. Considering a median depth of 84 million reads in the sequencing (Table 1), we converted junction counts to RP80M values (reads per 80 million mapped) using the total number of aligned reads across the autosomal chromosomes, which can be interpreted as the number of reads supporting the junction in an average library size (4).

**TABLE 1.** Associations of the splicing junctions with SCZ risk SNPs across the 13 brain regions.

| Tissue | N | Average mapped reads counts | ARL6IP4 |  | APOPT1 |  |  | CYP2D6 |  |
| --- | --- | --- | --- | --- | --- | --- | --- | --- | --- |
|  |  |  | rs1790121 |  | rs10431750 |  |  | rs133377 |  |
|  |  |  | Exon_2.3.1 | Exon_2.3.2 | Exon_1.4 | Exon_2.4 | Exon_3.4 | Exon_3.4 | Exon_2.4 |
| Amygdala | 88 | 83,912,256 | 1.45E-05 | 1.48E-10 | 0.0002 | 0.0165 | 0.021 | 0.0003 | 3.79E-06 |
| ACC | 109 | 88,136,513 | 3.16E-09 | 1.00E-13 | 2.39E-07 | 6.72E-05 | 2.40E-04 | 4.99E-08 | 5.34E-06 |
| Caudate | 144 | 86,167,404 | 4.26E-12 | 2.15E-26 | 1.01E-07 | 0.0046 | 9.52E-04 | 4.69E-11 | 8.47E-12 |
| Cerebellar | 125 | 89,274,842 | 1.38E-10 | 1.93E-25 | 0.0024 | 1.66E-04 | 0.0048 | 7.62E-13 | 9.09E-19 |
| Cerebellum | 154 | 82,622,299 | 5.23E-15 | 6.19E-30 | 0.0383 | 0.0092 | 7.67E-05 | 4.02E-10 | 1.47E-21 |
| Cortex | 136 | 82,272,249 | 1.05E-09 | 1.76E-26 | 0.0003 | 3.78E-05 | 2.76E-04 | 3.01E-07 | 5.20E-05 |
| DLPFC | 118 | 85,916,703 | 9.66E-15 | 2.85E-22 | 2.70E-05 | 3.86E-04 | 1.00E-04 | 2.65E-04 | 1.93E-06 |
| Hippocampus | 111 | 82,487,941 | 5.95E-09 | 2.64E-23 | 1.26E-06 | 0.0554 | 0.3127 | 3.02E-06 | 0.011 |
| Hypothalamus | 108 | 85,014,817 | 2.84E-11 | 3.45E-15 | 8.28E-04 | 0.011 | 5.07E-03 | 1.42E-08 | 6.22E-08 |
| Nucleus | 130 | 88,584,649 | 7.08E-10 | 6.76E-16 | 8.77E-11 | 0.0022 | 6.13E-05 | 2.38E-12 | 1.25E-13 |
| Putamen | 111 | 85,791,599 | 5.78E-05 | 4.97E-21 | 8.40E-04 | 0.8895 | 0.9736 | 3.72E-06 | 3.29E-06 |
| Spinal | 83 | 82,583,948 | 4.79E-08 | 4.11E-14 | 1.92E-04 | 0.0889 | 0.0568 | 0.0917 | 1.93E-04 |
| Sub | 80 | 80,973,047 | 0.0003 | 1.37E-14 | 0.0022 | 1 | 0.6795 | 3.35E-05 | 1.06E-04 |

False discovery rate (FDR) values are listed. ACC, Anterior cingulate cortex (BA24); Caudate, Caudate (basal ganglia); Cerebellar, Cerebellar Hemisphere; DLPFC, dorsolateral prefrontal cortex (BA9); Nucleus, Nucleus accumbens (basal ganglia); Putamen, Putamen (basal ganglia); Spinal, Spinal cord (cervical c-1); Sub, Substantia nigra.

### Genotyping data

Whole-genome sequencing (WGS) datasets were retrieved from dbGap upon authentication by the GTEx Consortium (Accession: phs000424.v7.p2). GTEx called all variants using GATK HaplotypeCaller (8). We extracted a total of 42,585,769 genomic variants which were then filtered step-by-step by using PLINK 1.9 (10) if they: 1) had a genotype missing rate >10% (272,734 variants); 2) had minor allele frequencies < 1% (31,110,395 variants); and 3) deviated from Hardy–Weinberg equilibrium (p-value < 1E-5, 791,170 variants). Finally, we retained 10,411,470 variants for further analysis.

### cis-acting eQTL analysis

cis-eQTL association was implemented separately by feature type (gene and junction) using Matrix eQTL R package (11) with the additive linear model, treating log2-transformed expression levels of each measurement (RPKM and RP80M) as the outcome. Low expression features (average counts < 0) were excluded before eQTL analysis. To control for potential confounding factors, we adjust for ancestry (first three principle components (PCs) from the genotype data) (12), sex, and the first K PCs of the normalized expression features, where K was calculated separately by feature type using the sva Bioconductor package (gene: 13 PCs, junction: 13 PCs). False discovery rate (FDR) was assessed using the Benjamini-Hochberg algorithm (BH) across all cis-eQTL tests within each chromosome. We considered all variant-gene pairs (eGene) and variant-junction pairs (eJunction) when the distance between features and SNP is less than 1MB.

### Co-localization of GWAS and eQTL associations

In order to assess the probability that molecular traits as estimated by cis-eQTLs and physiological traits as estimated by GWAS share the same causal variant, we co-localized 8,171,061 SCZ GWAS summary statistics (1) with our eGene and eJunction results. We used SMR and HEIDI tests for the co-localization analysis (13). We used the default parameters and performed for the genes and junctions. SNPs with LD r-squared between top-SNP > 0.90 or < 0.05 were excluded as well as one of each pair of the remaining SNPs with LD r-squared > 0.90. In addition, we conducted mappings of eGene and eJunction with SCZ GWAS separately. In the mapping process, any variants without either eQTL or GWAS association statistics were excluded.

### SNP annotation

ANNOVAR (14) was used for characterizing the categories of variants which include exonic, upstream, downstream, 3’-UTR, 5’-UTR, intronic, and intergenic regions. Roadmap/ENCODE2 chromatin-state signatures using a multivariate Hidden Markov Model (chromHMM) from brain tissue and cell types were extracted and visualized using WashU Epigenome Browser (http://epigenomegateway.wustl.edu/legac/). For the identification of the binding locations of transcription factors (TF), ENCODE TF binding data was downloaded (15). Then, we used BEDTools intersect (16) to match SNPs to ChIP-seq peaks. Brain histone ChIP-seq data were obtained from the ENCODE portal (https://www.encodeproject.org/). In addition, histone acetylation QTLs data (H3K9Ac ChIP-seq) generated from DLPFC of 433 individuals and DNA methylation QTLs data generated from 468 individuals were obtained from Brain xQTL Server (17).

### Functional enrichment

We used three tools (WebGestalt (18), DAVID (19), and gProfiler (20)) for overrepresentation enrichment analysis which help us identify biological pathways that are significantly enriched in a gene list. The transcript features were mapped to Entrez Gene IDs and subsequently to KEGG pathway. Gene ontologies (GO) biological process and GO molecular function (21) were also calculated. FDR (BH) and fold enrichment were imputed. FDR < 0.001 was used as threshold.

## Results

### Transcriptome-wide association study

Alterations in the brain have been demonstrated to underlie cognitive deficits associated with SCZ, including impairments in working memory and cognitive flexibility (22). To better understand the genetic interactions between expression features and genomic variations, we analyzed RNA-seq data of 13 brain regions from 1,497 postmortem brains in combination with WGS data. We comprehensively identify cis-QTLs and expression features (gene and exon-exon splicing junctions) in the human brain with quality-controlled SNP genotyping data from the same individuals using Matrix eQTL (11) in a genome-wide manner (see Methods for details). To conservatively define eGene and eJunction SNPs, we applied BH procedure for multiple testing implemented in Matrix eQTL to the p-values. To evaluate the extent of overlap between eQTL and GWAS signatures in SCZ and to identify putative causative genes from GWAS associations, we then performed co-localization analyses by SMR methods using default parameters. We also performed mapping for an in-depth investigation. After performing these procedures, we identified a total of 55 genes (SMR) and 89 genes (mapping) in eGene (Table S1 and Table S2) and 186 junctions within 78 genes (SMR) and 343 junctions within 133 genes in eJunction (mapping) (Table S3 and table S4) across the 13 brain regions (Figure 1 and Figure S1 and Figure S2).

**FIGURE 1.**
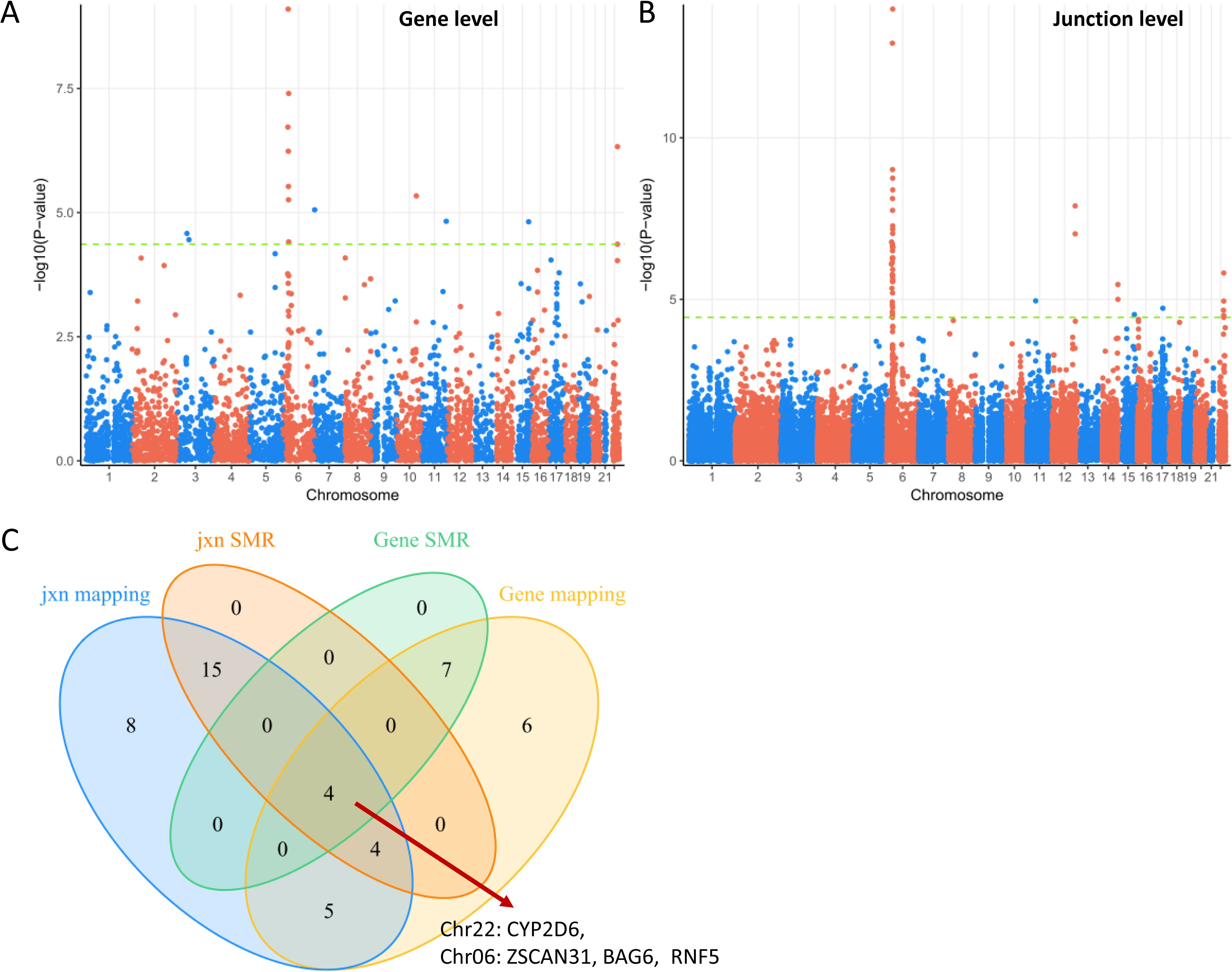
Identification of CYP2D6 as a top candidate for schizophrenia risk. Manhattan plot of DLPFC (Brodmann Area 9) in gene level (A) and junction level (B). (C) Venn Diagram of significant eGene and eJunction in dorsolateral prefrontal cortex (BA9). CYP2D6 was observed in all overlapped combinations. Jxn, exon-exon junction. Manhattan plots and Venn diagrams of other 12 brain regions are shown in Supplementary Figure S1-S3.

As expected, most of the signals are from the major histocompatibility complex (MHC) region: 32/55 (SMR) and 51/89 (mapping) genes within eGene (Table S1 and Figure 1A), and 41/78 and 74/133 (mapping) genes within eJunction (Table S1-4). The MHC is located in chromosome 6p21 and contains crucial regulators of the immune response. The MHC is the most gene dense and most polymorphic region of the human genome (23). Complement component 4 (C4) structural variation was recently demonstrated to be related to the expression of C4A and C4B in postmortem brain (24). In the current study, while C4A was detected in three brain regions in gene level by both SMR and mapping, C4A appeared across 10 brain regions in junction level by both SMR and mapping (Figure 1B and 1C, and Figure S3, Table S3 and Table S4). AS3MT was also replicated across several brain regions (Table S1-S4).

### Determination of splicing junctions

Alternative pre-mRNA splicing is a regulated process that results in a single gene coding for multiple proteins by including or excluding particular exons of a gene. This generates spliced mRNAs that direct the synthesis of a diverse set of proteins with varied biological functions. To illuminate our understanding of SCZ risk in splicing junction level, we systematically evaluated the exon-exon junctions in the human brain. We found eight splicing junctions located within four genes: Exon_2.3.1 and Exon_2.3.2 in *ARL6IP4*; Exon_1.4, Exon_2.4, and Exon_3.4 in *APOPT1*; Exon_3.4 and Exon_2.4 in *CYP2D6* (Figure 2 and Table 1). These junctions tag potential transcripts with alternative exonic boundaries (*ARL6IP4* Exon_2.3.1 and Exon_2.3.2) or exon skipping (*CYP2D6* Exon_3.4, *SNX19* Exon_8.10, and *APOPT1* Exon_1.4 and Exon_2.4) (Figure 2). In prior work, we have systematically characterized splice junctions between exon 8 and exon 10 in *SNX19* (7).

**FIGURE 2.**
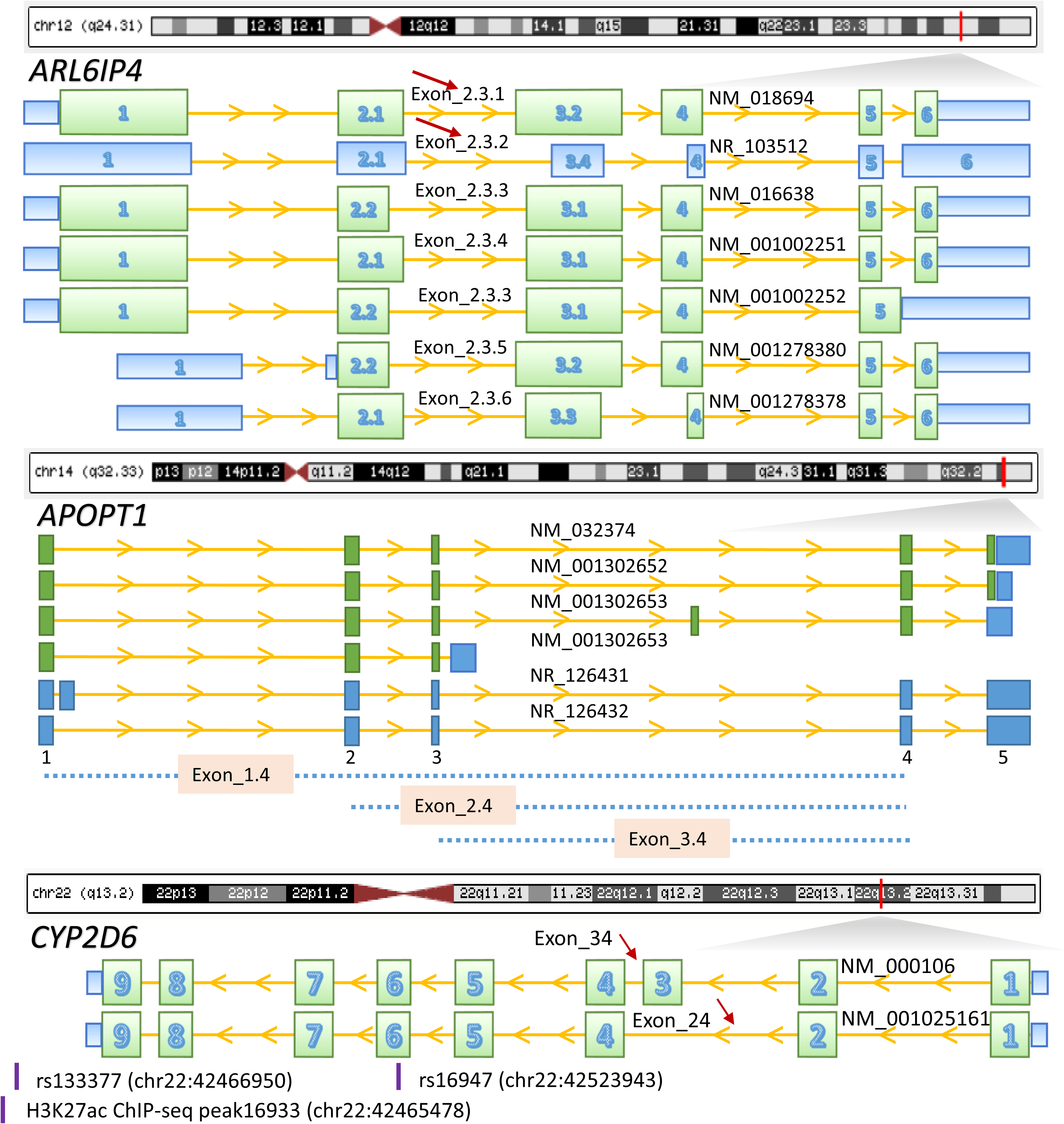
Splicing junctions on gene structures. Red bar at chromosomes indicate the gene physical position on the chromosome. Green boxes represent exons; blue boxes represent untranslated regions; solid yellow lines indicate introns; yellow arrows on yellow lines indicate gene transcriptional directions. Red arrows indicate identified junctions: Exon_2.3.1 (chr12:123465814-123466117) and Exon_2.3.2 (chr12:123465847-123466117) in *ARL6IP4*; Exon_1.4 (chr14:104029462-104053610), Exon_2.4 (chr14:104038158-104053610), and Exon_3.4 (chr14:104040508-104053610) in *APOPT1*; Exon_3.4 (chr22:42524947-42525034) and Exon_2.4 (chr22:42522995-42523448) in *CYP2D6*. The association of SNX19 splicing junction Exon_8.10 has been previously characterized in our prior work (7). Relative position of CYP2D6 SNPs and histone marker are indicated.

### Effects of Schizophrenia genetic risk on junctions

To determine how SCZ GWAS relates to feature expression, we next analyzed the trends of the associations. While SCZ risk allele is associated with down regulation of *APOPT1* junction Exon_3.4, all other risk alleles are associated with up regulation of junctions (Figure S4). Interestingly, SCZ risk allele is associate with up regulation of *APOPT1* junction Exon_2.4 (Figure 3). Opposing effects of SCZ risk allele on *APOPT1* Exon_3.4 and the other two junctions indicate their diverse functions. These robust effects are reproduced in the DLPFC and many other brain regions (Table 1). Based on the genomic recombination rate, the GWAS-eQTL loci are in a linkage block where recombination is not estimated to occur. In addition, a cluster of SNPs are in high LD with top schizophrenia GWAS-eQTL SNPs: *ARL6IP4* rs1790121, *APOPT1* rs10431750 and *CYP2D6* rs133377 (Figure S5).

**FIGURE 3.**
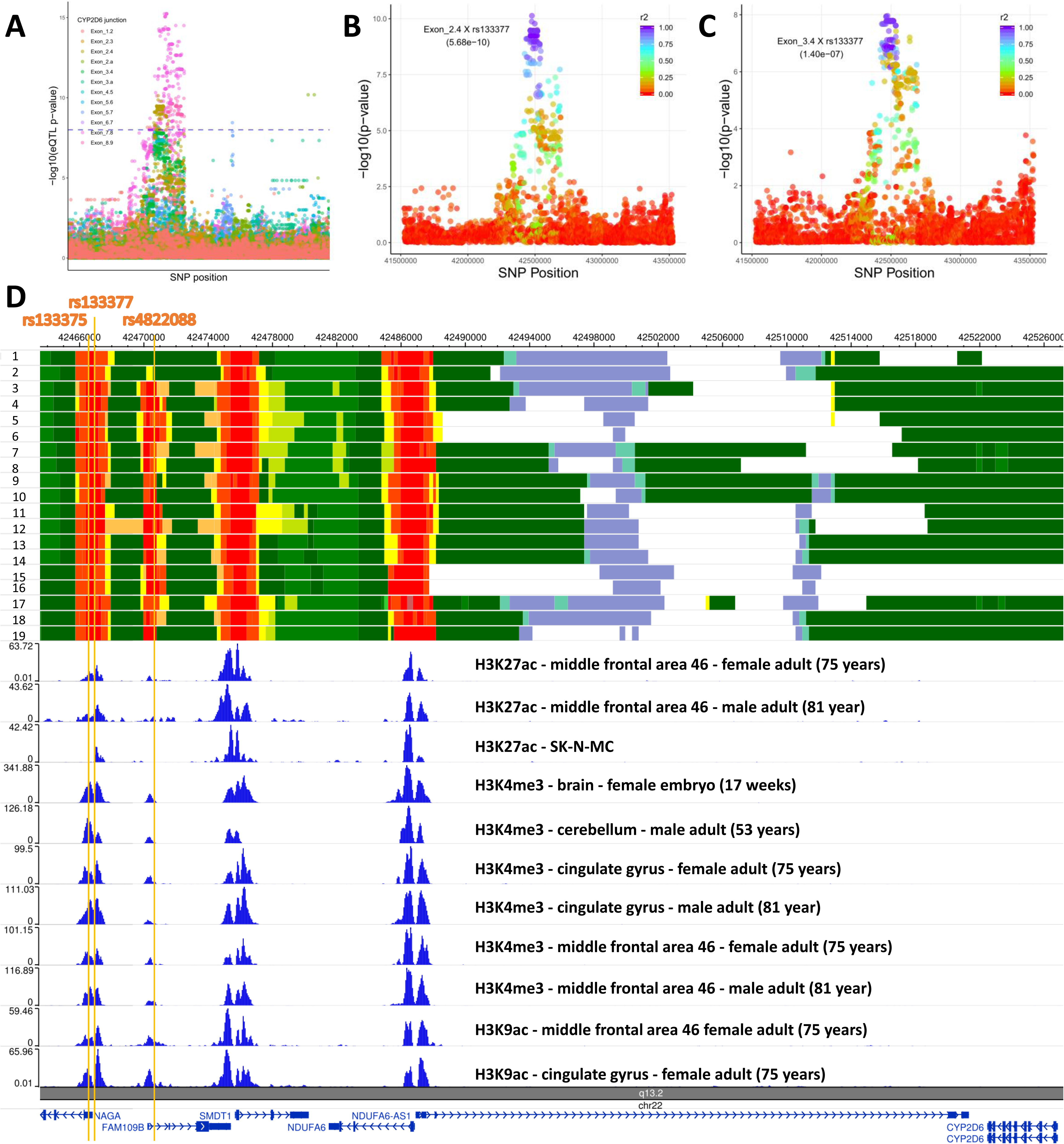
Identification of functional variants of CYP2D6. Association of SNP-junction pairs (eJunction) of CYP2D6. Association of junctions Exon_2.4 (B) and Exon_3.4 (C) with SNPs upstream and downstream of rs133377. *r*^2^ was estimated using DLPFC samples. See association results from other 12 brain regions in Supplementary Figure S6-S8. (D) Schizophrenia splicing SNPs are located within active transcription start sites. Upper section: ENCODE chromatin activation states in human brain from Ganglion eminence derived primary cultured neurospheres (1), cortex derived primary cultured neurospheres (2), anterior caudate (3, 4), cingulate gyrus (5, 6), hippocampus middle (7, 8), inferior temporal lobe (9, 10), dorsolateral prefrontal cortex (11, 12), substantia nigra (13, 14), NH-A astrocytes primary cells (15, 16), germinal matrix (17), angular gyrus (18, 19). See details in Supplementary Table S6. Middle section: ChIP-seq of H3K27ac, H3K4me3, and H3K9ac. Lower section: reference genes. Note: yellow bars represent coordinates of rs133375, rs133377 and rs4822088.

Strikingly, we found *CYP2D6* to be the top gene to be determined in both eGene and ejunction across the 13 brain regions using both SMR and mapping (Figure 1, Figure S1-S3, Table S1-S4). Therefore, we further investigated the *CYP2D6* region. While multiple junctions were determined to be significant at the junction level across the 13 brain regions, we also found Exon_2.4, Exon_3.4, and Exon_7.8 to be significantly associated with SCZ risk SNPs (Figure 4A, Figure S6-S8 and Table S5). Interestingly, splicing occurred between exon 2 and exon 4 which resulted in maintaining or skipping exon 3. Exon_78 was included in all *CYP2D6* transcripts which represents gene level (Figure 2).

**FIGURE 4.**
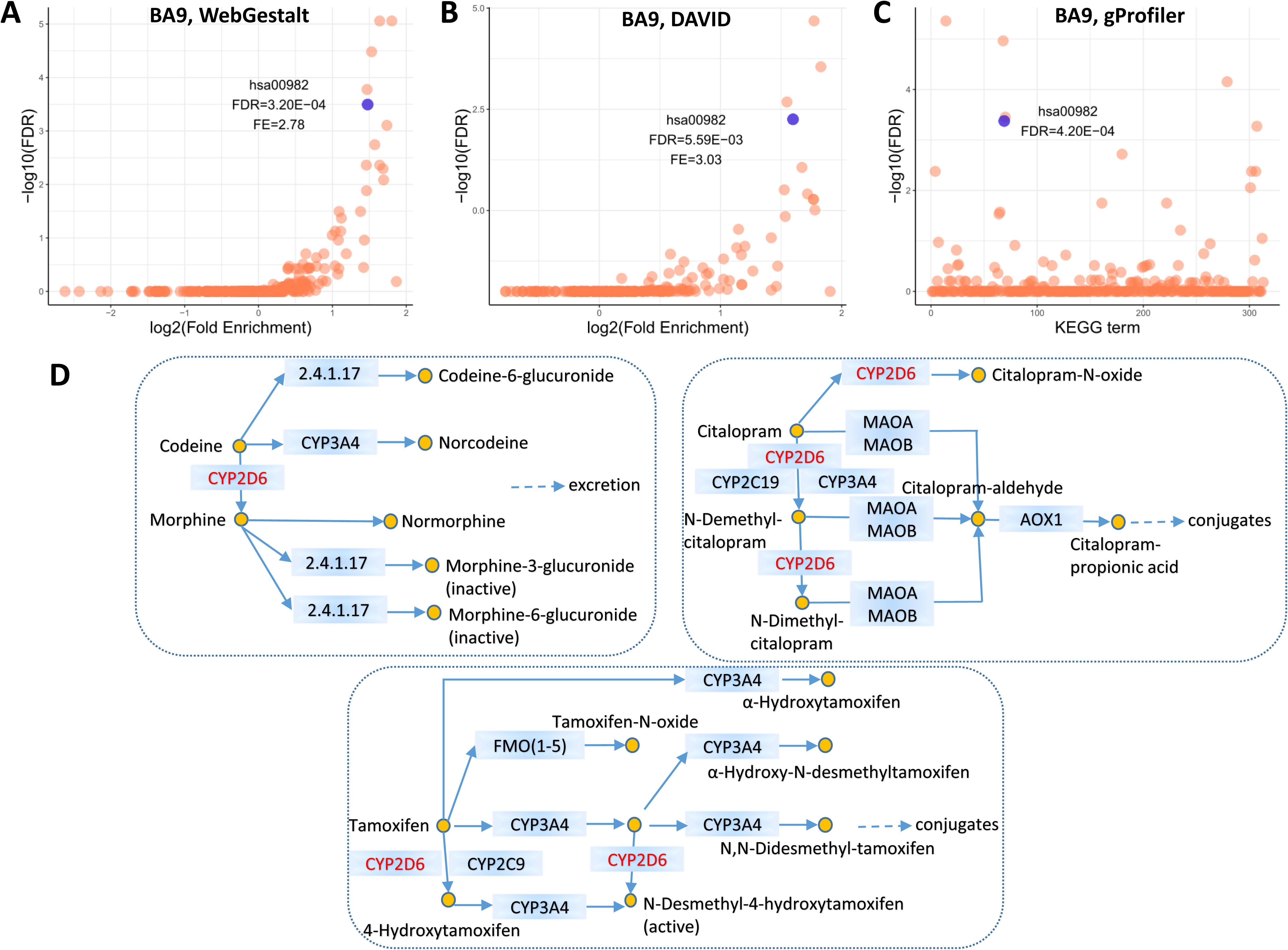
Pathway analysis of overlapped significant genes of eJunctions and eGenes. KEGG pathways enriched for 798 genes that are both significant in gene level and junction level in dorsolateral prefrontal cortex (BA9) by using WebGestalt (A), DAVID (B), and gProfiler (C). Blue dots are KEGG pathway has00982 in which CYP2D6 is involved. FDR: false discovery rate. Results of other 12 brain regions are shown in Supplementary Figure S11 (WebGestalt), Figure S12 (DAVID), and Figure S13 (gProfiler). (D) CYP2D6 in the drug metabolism process of KEGG pathway hsa00982. 2.4.1.17: UGT1A(1,3-10), UGT2A(1-3), UGT2B(4,7,10,11,17,28).

### Functional characterization of eJunction SNPs in SCZ

Determining the underlying causal variants of complex disorders can be challenging because of the complex linkage disequilibrium patterns between SNPs. We next attempted to functionally characterize eQTL SNPs by classifying the 255 SNPs in SCZ GWAS eJunction according to the definition in ANNOVAR (14). As expected, most of the SNPs are non-exonic which accounts for 85.25% in SCZ GWAS eGene and 87.09% in SCZ GWAS eJunction (Figure S9 and Table S5). There are a total of seven exonic single nucleotide variant (SNV), four of which are non-synonymous. Only one SNV was found in our target genes, which is, rs16947, located within exon 6 of *CYP2D6* gene (NM_000106) (Figure 2 and Table S5). Our CYP2D6 regional (2MB, about 5,000 SNPs) LD analysis showed that SNPs in high LD with rs16947 are strongly associated with the abundance of splicing junctions (Exon_34 and Exon_24) in DLPFC and other brain regions (Table S5).

Enhancers have emerged as key cis-regulatory elements that play important regulatory roles in gene transcription, and they often reside distally from their target of regulation (25). Using ENCODE chromatin activation states from brain, we found that three SCZ splicing QTLs (rs133375, rs133377 and rs4822088) are located within active transcription start sites (TSS) (Figure 4D). Of the three SCZ splicing SNPs, rs133377 showed to be intercrossed with active transcription state across all the brains (Table S6). Three histone marks on H3K27ac, H3K4me3, and H3K9ac have been established to be associated with active transcription (26, 27). As expected, the three SNPs we identified are located within the chromatin modification peaks (Figure 4D). Note that the three SNPs are in high LD with rs16947 (*r*^2^ = 0.9539) (Figure S10 and Table S5). In addition, we analyzed another independent histone acetylation data and found rs133377 is significantly associated with a histone acetylation site, peak16933 in a 2 MB sliding window (chr22:42465478, p-value = 3.78E-8) (Figure S11 and Table S7). DNA methylation plays important roles in epigenetically regulating gene expression in the brain. Using DNA methylation data of postmortem DLPFC brain from 468 individuals (17), we found 27 significant SNP-CpG pairs scattered around *CYP2D6* (Table S7), implying they function together to regulate gene transcription.

### Enrichment analysis of eJunction

To further verify our systematic genomics results and gain mechanistic insight, we next took an unbiased data-driven approach and performed a comprehensive gene-set pathway enrichment analysis to explore the potential functional implications of the genome-wide significant genes overlapped in junction level and gene level using WebGestalt, DAVID and gProfiler across the 13 brain regions. An average of 822 genes were determined and used for the enrichment analysis, and more than 300 pathways were imputed by the three tools. Two top pathways that include CYP2D6 are associated with metabolism of xenobiotics by cytochrome P450 (hsa00980) and drug metabolism (hsa00982) and were consistently identified by each of the three tools across the 13 brain regions (Figure 4A, 4B, 4C, Figure S12-S14, and Table S8-S10). When we took an in-depth examination of the KEGG hsa00982 pathway, we found that CYP2D6, together with several other determined genes, were associated with the metabolic process of psychiatric drugs: tamoxifen, codeine and morphine, and citalopram (Figure 4D and Figure S12-S14). Similarly, GO biological process analyses showed *CYP2D6* and related genes play a role in steroid metabolic process in DLPFC and other brain regions (GO:0008202) (Table S11). GO molecular function analyses showed CYP2D6 and 13 related genes involved in drug binding (GO:0008144) in DLPFC and other brain regions (Table S12).

## Discussion

In this study, we analyzed a large-scale transcriptome derived from 1,497 human postmortem brain samples in combination with WGS data. SCZ is a complex genetic disorder and over a hundred loci have been identified. However, it remains unclear how genetics affect the abundance of expression features. We identified expression features and potential effective splicing transcripts underlying SCZ susceptibility using the most recent GTEx brain RNA-seq data (released on June 30, 2017) (8) and 2018 SCZ GWAS summary statistics (1). We identify *CYP2D6* to be an important candidate gene through several unbiased strategies: 1) by comparing gene overlapping using SMR and mapping in gene and junction level across 13 brain regions; and 2) through hypothesis-free over-representation enrichment analysis of eJunction genes. Furthermore, by analyzing multiple sources of ChIP-seq data generated from postmortem brains, we identify causal SNPs associated with CYP2D6.

Our principal findings include four major observations. First, we have identified a total of 171 genes across 13 brain regions for which genetic variation for expression co-localize with genetic variation for SCZ risk. Some differences in results are expected using alternative co-localization methods and references. Using 206 postmortem brain DLPFC samples from normal individuals, Takata et al evaluated SCZ risk loci that are involved in splicing events based on assembled transcripts (3). In the current study, we replicated reported genes, for example, *SNX19, ARL6IP4, APOPT1* (3), *C4A* and *C4B* (24), and discovered new genes such as *CYP2D6*.

Second, we identified a list of exon-exon junctions that tag genetic transcripts. In prior work, most analyses focus almost exclusively on the marginal eQTL signal which typically represents the primary, or most significant, eQTL signal, rather than dissecting this signal into multiple independent features for each gene. While predicted transcripts are more likely to be used as a reference, we used exon-exon junctions to tag specific transcripts which considerably increases specificity. Our eJunction analyses identify eight splicing junctions that are located in four genes. Our recent study has systematically evaluated SNX19 Exon_8.10 transcripts and their potential susceptibility to SCZ (7). APOPT1 was demonstrated to be located in mitochondria matrix, and involved in possessing an N-terminal mitochondrial targeting signal (28). ARL6IP4 play a functional role in pre-mRNA splicing (29, 30). However, the mechanisms of APOPT1 and ARL6IP4 are still largely unknown.

Third, we illustrated an underlying mechanism for SCZ risk. There is accumulating evidence that Cytochrome P450 2D6 (CYP2D6) is involved in the metabolism of clinically used drugs (31). A clinical study reported that CYP2D6 plays an important role in controlling the state of aripiprazole in the plasma which has been established as a form of treatment for SCZ (32). CYP2D6 has also been reported to have significant effects on psychotic symptoms (33) and cognitive performance (34) of SCZ patients following risperidone treatment. Our data-driven enrichment analysis identified CYP2D6 as a significant eGene and eJunction gene and as a key component of the drug metabolism pathway (specifically related to codeine, citalopram, morphine and tamoxifen). CYP2D6/codeine has been highlighted as an antidepressant gene/drug pair in clinical therapy by Clinical Pharmacogenetics Implementation Consortium (35). Codeine is bioactivated into morphine by CYP2D6 to exert its analgesic effect. Morphine, a strong opioid agonist, acts directly on the central nervous system and has been strongly implicated in addiction and SCZ pathophysiology (36). Citalopram is a selective serotonin reuptake inhibitor used to treat major depression disorders (37). Tamoxifen is well-known as an estrogen receptor modulator commonly used to treat breast cancer. In addition, we observed CYP2D6 to be involved in the metabolism of neuroactive steroids which are present in human post-mortem brain tissue. In fact, concentrations of neuroactive steroids are known to be altered in subjects with SCZ and bipolar disorder (38). Overall, these findings highlight CYP2D6 as an important candidate for further biological investigation. For example, it would be interesting to investigate how CYP2D6 interacts with the top identified SCZ risk genes and contributes to SCZ risk. It would also be interesting to determine how the diverse CYP2D6 transcripts differentially affect the metabolism of psychiatric drugs.

Finally, by characterizing the properties of the detected eJunction SNPs, we found one non-synonymous SNV, rs16947, located within CYP2D6 exon 6, to be associated with abundance of splicing junctions related to its exon 3 skipping. This leads to an in-frame deletion that shortens the translated protein by 51 amino acids. This risk allele of the SNV could result in an amino acid substitution that has been shown to reduce enzyme activity after recombinant cDNA transfection (39, 40). On the other hand, we found three SNPs, which are in strong LD with rs16947, to be located within active chromatin modification regions in the brain, indicating that they are actively involved in direct regulation of CYP2D6 gene transcription (26, 27). Thus, taken together, it is conceivable these alterations (changes in protein structure and/or chromatin modifications) affect the transcriptional activity of *CYP2D6*, leading to downstream changes in gene expression, which could further alter enzyme activity in drug metabolism. This alternative pre-mRNA splicing may also contribute to the extensive variability in CYP2D6 activity observed across individuals (41).

Although the sample size in this study is substantial (N=1,497), the sample size of each brain region is relatively small (mean = 115), so increasing sample size could help identify additional brain functional SNPs and splicing junctions with increased confidence. Along the same lines, quantifying feature differences between SCZ and normal controls samples would also be informative. While we provide new targets implicated in SCZ, further understanding the regulatory mechanisms of the genomic markers and gene structures identified here in animal or cellular models (for example, human derived neurons) would be of great value.

In summary, we comprehensively analyzed expression features that are mediated by genomic markers across the human brain regions, described the characteristics of these SNPs, and demonstrated that the list of brain SNPs can be used to identify plausible candidate transcripts/variants that are causally associated with SCZ. These findings will be of great use in generating new animal and cellular models for SCZ.

## Acknowledgments and disclosures

We thank all the staff especially John B. Hanks at the Stanford Research Computing Center for their excellent computational support. We thank Anna Shcherbina at Stanford University, Yuxing Liao at Baylor College of Medicine, and Hedi Peterson at University of Tartu for their suggestions and help. This work was supported by grants from the Stanford University School of Medicine and a Siebel Fellowship awarded to S.C.. Data were generated by the GTEx Project supported by the Common Fund of the National Institutes of Health.

## Author contributions

L.M. conceived the study, conducted the analysis, and wrote the draft. L.M. and S.C. collected the data. S.C. provided intellectual input and financial support for the study. All authors contributed to the editing of the manuscript, and approved the final manuscript.

## Supplementary Table

Supplementary Table S1. SNPs that are significant in schizophrenia GWAS are significantly associated with genes in 13 brain regions by SMR & HEIDI methods

Supplementary Table S2. SNPs that are significant in schizophrenia GWAS are significantly associated with genes in 13 brain regions by mapping

Supplementary Table S3. SNPs that are significant in schizophrenia GWAS are significantly associated with junctions in 13 brain regions by SMR & HEIDI methods

Supplementary Table S4. SNPs that are significant in schizophrenia GWAS are significantly associated with junctions in 13 brain regions by mapping

Supplementary Table S5. Schizophrenia risk SNPs are significantly associated with CYP2D6 junctions across 13 brain regions

Supplementary Table S6. Samples and chromatin state of CYP2D6 SNPs shown in Figure 4D

Supplementary Table S7. Association of CYP2D6 schizophrenia risk eJunction SNPs with histone acetylation peaks and CpG sites

Supplementary Table S8. KEGG pathway of overlapped genes regulated by eGene and eJunction SNPs across the 13 brain regions by WebGestalt

Supplementary Table S9. KEGG pathway of overlapped genes regulated by eGene and eJunction SNPs across the 13 brain regions by DAVID

Supplementary Table S10. KEGG pathway of overlapped genes regulated by eGene and eJunction SNPs across the 13 brain regions by gProfiler

Supplementary Table S11. Steroid metabolic process of gene ontologies (GO:0008202) across multiple brain regions by DAVID

Supplementary Table S12. Drug binding of gene ontologies (GO:0008144) across multiple brain regions by DAVID

## Supplementary Figure

Supplementary Figure S1. Manhattan plot of the 12 brain regions in gene level. Manhattan plots of dorsolateral prefrontal cortex (BA9) are shown in Figure 1.

Supplementary Figure S2. Manhattan plot of the 12 brain regions in junction level. Manhattan plots of dorsolateral prefrontal cortex (BA9) are shown in Figure 1.

Supplementary Figure S3. Venn diagram of eGene and eJunction analysis across the 12 brain regions. Manhattan plots of dorsolateral prefrontal cortex (BA9) are shown in Figure 1.

Supplementary Figure S4. Schizophrenia genetic risk effect on exon-exon junctions. Association of SNP-junction pairs: rs1790121 - ARL6IP4 Exon_2.3.1 (up) and Exon_2.3.2 (up); rs10431750 - APOPT1 Exon_1.4 (up) and Exon_2.4 (up) and Exon_3.4 (down); rs133377 - CYP2D6 Exon_34 (up) and Exon_24 (up). The junction abundance is from DLPFC. Red arrow indicates schizophrenia risk allele is associated with up regulation of junction, while blue arrow is associated with downregulation. SNX19 splicing junction Exon_8.10 was shown in our recent paper (Ma, et al., 2019).

Supplementary Figure S5. Regional schizophrenia GWAS signature co-localized with our identified variants that influence splicing of APOPT1 (Exon_1.4 and Exon_2.4 and Exon_3.4), CYP2D6 (Exon_34 and Exon_24). Linkage disequilibrium is colored with respect to ARL6IP4 rs1790121, APOPT1 rs10431750, and CYP2D6 rs133377. See SNX19 in our prior work (Ma, et al., 2019).

Supplementary Figure S6. Association of SNP-junction pairs (eJunction) of CYP2D6. See association results from DLPFC regions in Figure 4A.

Supplementary Figure S7. Association of junctions Exon_2.4 with SNPs upstream and downstream of rs133377. r2 was estimated using corresponding brain regions. See association results from DLPFC regions in Figure 4B.

Supplementary Figure S8. Association of junctions Exon_3.4 with SNPs upstream and downstream of rs133377. r2 was estimated using corresponding brain regions. See association results from DLPFC regions in Figure 4C.

Supplementary Figure S9. Characterization of identified schizophrenia GWAS eJunction SNPs. Pie charts indicating proportions of SNPs annotated with each functional category (exonic, upstream, downstream, 3’-UTR, 5’-UTR, splicing, intronic and intergenic).

Supplementary Figure S10. Linkage disequilibrium (LD) plot of identified schizophrenia GWAS functional SNPs. r2 was estimated using DLPFC data. See LD r2 at Supplementary Table S5.

Supplementary Figure S11. Association of rs133377 (chr22:42466950) with 70 histone acetylation peaks in 2 MB region around CYP2D6.

Supplementary Figure S12. Pathway analysis of overlapped significant genes of eJunctions and eGenes by WebGestalt across 12 brain regions.

Supplementary Figure S13. Pathway analysis of overlapped significant genes of eJunctions and eGenes by DAVID across 12 brain regions.

Supplementary Figure S14. Pathway analysis of overlapped significant genes of eJunctions and eGenes by gProfiler across 12 brain regions.

## References

1. Pardinas AF, Holmans P, Pocklington AJ,xs et al: Common schizophrenia alleles are enriched in mutation-intolerant genes and in regions under strong background selection. Nat Genet 2018; 50:381–389

2. Schizophrenia_Working_Group_of_the_Psychiatric_Genomics_Consortium: Biological insights from 108 schizophrenia-associated genetic loci. Nature 2014; 511:421–427

3. Takata A, Matsumoto N, Kato T: Genome-wide identification of splicing QTLs in the human brain and their enrichment among schizophrenia-associated loci. Nat Commun 2017; 8:14519

4. Jaffe AE, Straub RE, Shin JH, et al: Developmental and genetic regulation of the human cortex transcriptome illuminate schizophrenia pathogenesis. Nat Neurosci 2018; 21:1117–1125

5. Steijger T, Abril JF, Engstrom PG, et al: Assessment of transcript reconstruction methods for RNA-seq. Nat Methods 2013; 10:1177–1184

6. Li M, Jaffe AE, Straub RE, et al: A human-specific AS3MT isoform and BORCS7 are molecular risk factors in the 10q24.32 schizophrenia-associated locus. Nat Med 2016; 22:649–656

7. Ma L, Semick SA, Chen Q, et al: Schizophrenia risk variants influence multiple classes of transcripts of sorting nexin 19 (SNX19). Mol Psychiatry 2019;

8. GTEx_Consortium, Laboratory DA, Coordinating Center -Analysis Working G, et al: Genetic effects on gene expression across human tissues. Nature 2017; 550:204–213

9. Harrow J, Frankish A, Gonzalez JM, et al: GENCODE: the reference human genome annotation for The ENCODE Project. Genome Res 2012; 22:1760–1774

10. Chang CC, Chow CC, Tellier LC, et al: Second-generation PLINK: rising to the challenge of larger and richer datasets. Gigascience 2015; 4:7

11. Shabalin AA: Matrix eQTL: ultra fast eQTL analysis via large matrix operations. Bioinformatics 2012; 28:1353–1358

12. Price AL, Patterson NJ, Plenge RM, et al: Principal components analysis corrects for stratification in genome-wide association studies. Nat Genet 2006; 38:904–909

13. Zhu Z, Zhang F, Hu H, et al: Integration of summary data from GWAS and eQTL studies predicts complex trait gene targets. Nat Genet 2016; 48:481–487

14. Wang K, Li M, Hakonarson H: ANNOVAR: functional annotation of genetic variants from high-throughput sequencing data. Nucleic Acids Res 2010; 38:e164

15. Kheradpour P, Kellis M: Systematic discovery and characterization of regulatory motifs in ENCODE TF binding experiments. Nucleic Acids Res 2014; 42:2976–2987

16. Quinlan AR, Hall IM: BEDTools: a flexible suite of utilities for comparing genomic features. Bioinformatics 2010; 26:841–842

17. Ng B, White CC, Klein HU, et al: An xQTL map integrates the genetic architecture of the human brain’s transcriptome and epigenome. Nat Neurosci 2017; 20:1418–1426

18. Wang J, Vasaikar S, Shi Z, et al: WebGestalt 2017: a more comprehensive, powerful, flexible and interactive gene set enrichment analysis toolkit. Nucleic Acids Res 2017; 45:W130–W137

19. Huang da W, Sherman BT, Lempicki RA: Systematic and integrative analysis of large gene lists using DAVID bioinformatics resources. Nat Protoc 2009; 4:44–57

20. Reimand J, Arak T, Vilo J: g:Profiler--a web server for functional interpretation of gene lists (2011 update). Nucleic Acids Res 2011; 39:W307–315

21. Gene_Ontology_Consortium: Gene Ontology Consortium: going forward. Nucleic Acids Res 2015; 43:D1049–1056

22. Weinberger DR, Berman KF, Zec RF: Physiologic dysfunction of dorsolateral prefrontal cortex in schizophrenia. I. Regional cerebral blood flow evidence. Arch Gen Psychiatry 1986; 43:114–124

23. Trowsdale J, Knight JC: Major histocompatibility complex genomics and human disease. Annu Rev Genomics Hum Genet 2013; 14:301–323

24. Sekar A, Bialas AR, de Rivera H, et al: Schizophrenia risk from complex variation of complement component 4. Nature 2016; 530:177–183

25. Heintzman ND, Ren B: Finding distal regulatory elements in the human genome. Curr Opin Genet Dev 2009; 19:541–549

26. Creyghton MP, Cheng AW, Welstead GG, et al: Histone H3K27ac separates active from poised enhancers and predicts developmental state. Proc Natl Acad Sci U S A 2010; 107:21931–21936

27. Gates LA, Shi J, Rohira AD, et al: Acetylation on histone H3 lysine 9 mediates a switch from transcription initiation to elongation. J Biol Chem 2017; 292:14456–14472

28. Melchionda L, Haack TB, Hardy S, et al: Mutations in APOPT1, encoding a mitochondrial protein, cause cavitating leukoencephalopathy with cytochrome c oxidase deficiency. Am J Hum Genet 2014; 95:315–325

29. Sasahara K, Yamaoka T, Moritani M, et al: Molecular cloning and expression analysis of a putative nuclear protein, SR-25. Biochem Biophys Res Commun 2000; 269:444–450

30. Li Q, Zhao H, Jiang L, et al: An SR-protein induced by HSVI binding to cells functioning as a splicing inhibitor of viral pre-mRNA. J Mol Biol 2002; 316:887–894

31. Phillips KA, Veenstra DL, Oren E, et al: Potential role of pharmacogenomics in reducing adverse drug reactions: a systematic review. JAMA 2001; 286:2270–2279

32. Suzuki T, Mihara K, Nakamura A, et al: Effects of genetic polymorphisms of CYP2D6, CYP3A5, and ABCB1 on the steady-state plasma concentrations of aripiprazole and its active metabolite, dehydroaripiprazole, in Japanese patients with schizophrenia. Ther Drug Monit 2014; 36:651–655

33. Bartecek R, Jurica J, Zrustova J, et al: Relevance of CYP2D6 variability in first-episode schizophrenia patients treated with risperidone. Neuro Endocrinol Lett 2012; 33:236–244

34. Zeng L, Kang C, Yuan J, et al: CYP2D6 polymorphisms are associated with effects of risperidone on neurocognitive performance in schizophrenia. Schizophr Res 2017; 188:50–51

35. Crews KR, Gaedigk A, Dunnenberger HM, et al: Clinical Pharmacogenetics Implementation Consortium (CPIC) guidelines for codeine therapy in the context of cytochrome P450 2D6 (CYP2D6) genotype. Clin Pharmacol Ther 2012; 91:321–326

36. Stefano GB, Kralickova M, Ptacek R, et al: Low dose morphine adjuvant therapy for enhanced efficacy of antipsychotic drug action: potential involvement of endogenous morphine in the pathophysiology of schizophrenia. Med Sci Monit 2012; 18:HY23–26

37. Cipriani A, Purgato M, Furukawa TA, et al: Citalopram versus other anti-depressive agents for depression. Cochrane Database Syst Rev 2012:CD006534

38. Marx CE, Stevens RD, Shampine LJ, et al: Neuroactive steroids are altered in schizophrenia and bipolar disorder: relevance to pathophysiology and therapeutics. Neuropsychopharmacology 2006; 31:1249–1263

39. Marcucci KA, Pearce RE, Crespi C, et al: Characterization of cytochrome P450 2D6.1 (CYP2D6.1), CYP2D6.2, and CYP2D6.17 activities toward model CYP2D6 substrates dextromethorphan, bufuralol, and debrisoquine. Drug Metab Dispos 2002; 30:595–601

40. Yu A, Kneller BM, Rettie AE, et al: Expression, purification, biochemical characterization, and comparative function of human cytochrome P450 2D6.1, 2D6.2, 2D6.10, and 2D6.17 allelic isoforms. J Pharmacol Exp Ther 2002; 303:1291–1300

41. Zanger UM, Raimundo S, Eichelbaum M: Cytochrome P450 2D6: overview and update on pharmacology, genetics, biochemistry. Naunyn Schmiedebergs Arch Pharmacol 2004; 369:23–37

